# TRIAC disrupts cerebral thyroid hormone action via a negative feedback loop and heterogenous distribution among organs

**DOI:** 10.1101/2022.08.12.502299

**Authors:** Ichiro Yamauchi, Takuro Hakata, Yohei Ueda, Taku Sugawa, Ryo Omagari, Yasuo Teramoto, Shoji F Nakayama, Daisuke Nakajima, Takuya Kubo, Nobuya Inagaki

## Abstract

3,3’,5-triiodothyroacetic acid (TRIAC) is a metabolite of endogenous thyroid hormones (THs) that can bind to and activate TH receptors. As TRIAC was previously detected in sewage effluent, we aimed to investigate exogenous TRIAC’s potential for endocrine disruption. We administered either TRIAC or 3,3’,5-triiodo-L-thyronine (LT3) to both euthyroid mice and 6-propyl-2-thiouracil-induced hypothyroid mice. In hypothyroid mice, TRIAC alleviated growth retardation, suppressed the hypothalamus-pituitary-thyroid (HPT) axis, and upregulated TH-responsive genes in the pituitary gland, liver, and heart. We observed that, unlike LT3, TRIAC does not upregulate the expression of TH-responsive genes in the cerebrum. Measurement of organ-specific TRIAC levels suggested that TRIAC was not efficiently trafficked into the cerebrum. Furthermore, by analyzing euthyroid mice, we found that cerebral TRIAC levels did not increase despite TRIAC administration at higher concentrations, whereas serum and cerebral TH levels were substantially decreased. Hence, TH-responsive genes in the cerebrum appear to be downregulated by TRIAC. In summary, TRIAC administration decreases circulating TH levels by suppressing the HPT axis, while the consequent attenuation of TH actions was compensated by TRIAC in peripheral tissues but not in the cerebrum due to the relative impermeability of the blood–brain barrier towards TRIAC. We verified that exogenous TRIAC disrupts TH actions in the cerebrum. This disruption is apparently due to the additive effects of circulating endogenous THs being depleted via a negative feedback loop involving the HPT axis and heterogenous distribution of TRIAC among different organs. Our findings indicate that environmental TRIAC poses a potential neurodevelopmental risk.

## Introduction

Thyroid hormones (THs) are essential for key biological processes, including development, growth, differentiation, and homeostasis. In many species, the hypothalamus-pituitary-thyroid (HPT) axis plays a substantial role in regulating TH action (1). In brief, thyroid stimulating hormone (TSH) secreted from the pituitary gland promotes the synthesis and secretion of THs by the thyroid gland. TSH production is stimulated by thyrotropin-releasing hormone (TRH) from the hypothalamus. THs inhibit TSH and TRH secretion via a negative feedback loop. In addition to the HPT axis, various components are necessary for TH action, including transporters, iodothyronine deiodinases (DIO1, DIO2, and DIO3), thyroid hormone receptors (THRs), and clearance pathways such as glucuronidation.

The two major THs that bind to and activate THRs are 3,3’,5,5’-tetraiodothyronine (T4) and 3,3’,5-triiodothyronine (T3). T3 is considered the active form because it is more potent than T4 at activating THRs. T3 is secreted from the thyroid gland along with T4 but most circulating T3 is the product of T4 deiodination by DIO1 and DIO2 (2). Meanwhile, some TH metabolites can be detected in human sera, including 3,3’-diiodothyronine, 3,3’,5’-triiodothyronine, 3,3’,5-triiodothyroacetic acid (TRIAC, also known as TA3), and 3,3’,5,5’-tetraiodothyroacetic acid (3). Among these metabolites, TRIAC is well known for its robust biological activity. TRIAC has a similar affinity to T3 for THRα1 and greater affinity than T3 for THRβ1 and THRβ2 (4–6). TRIAC exerts its effects on the HPT axis as well as other organs, including the heart, bones, and liver of animals as reviewed elsewhere (7).

Endocrine-disrupting chemicals (EDCs) are exogenous chemicals that interfere with the normal function of hormones (8). EDCs pose an adverse risk to human health while also impacting natural ecosystems and wildlife. Since people may be exposed to EDCs throughout their entire life, the management of environmental EDCs is crucial (9). EDCs can disrupt TH actions through a variety of mechanisms. For example, bisphenol A disrupts by acting on THRs (10), a TH transporter MCT8 (11), and DIO3 (12). A second example is the lowering of circulating TH levels by polychlorinated biphenyls, polybrominated diphenyl ethers, as well as phthalates (1). Perchlorate inhibition of iodide uptake by the thyroid gland is a third example of such disruption (13). Thus, the disruption mechanisms of EDCs are both varied and complex, and must be determined for each EDC.

Our previous work highlighted TRIAC’s potential to act as an EDC. THR agonist activity in sewage effluent has been determined in multiple countries (14–17). We determined TRIAC is a main contributor of THR agonist activity in sewage effluents (18). Importantly, sewage treatment plants can release chemicals into environmental water, which can subsequently be consumed by humans as well as wildlife. We reemphasize that TRIAC binds to THRs as a ligand and exhibits agonist activity equal to or greater than that of T3 (4–7). The potential of exogenous TRIAC for endocrine disruption should be urgently investigated to determine its risk to human health.

The aim of the study presented here was to elucidate the impact of exogenous TRIAC on TH action. We administered TRIAC to mice and analyzed the animals using our methods to evaluate various elements that regulate TH action (19–21). To determine TRIAC’s effects, we administered 3,3’,5-triiodo-L-thyronine (LT3) to other mice for comparison. We thereby determined that exogenous TRIAC appears to attenuate TH actions exclusively in the cerebrum.

## Results

### TRIAC administration to hypothyroid mice

THs are essential to skeletal growth as evidenced by the phenotype of THRβ knockout mice (22). Since the growth rate of C57BL/6 mice slows considerably after six weeks of age (23), we determined the effects of THs on growth and other processes by administering LT3 and TRIAC beginning at three weeks of age immediately after weaning (Fig. 1). We administered LT3 and TRIAC to euthyroid mice without impairment of TH secretion. Although we administered 0.1 ppm of either LT3 and TRIAC as a supplemental dose or 1 ppm as a high dose to the mice, we did not observe significant differences in growth curves based on body weight and naso-anal length except for the naso-anal length of mice administered with 1 ppm TRIAC (Supplementary Fig. S1).

**Fig. 1:**
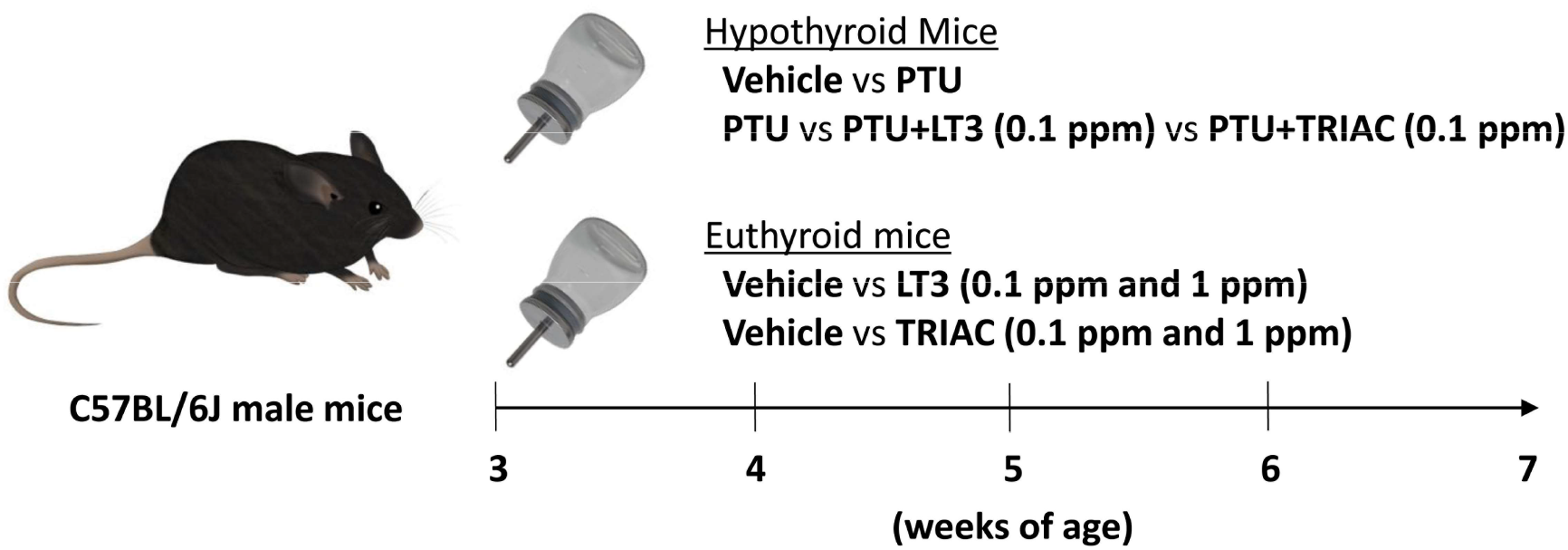
Schema for generating mouse cohorts. “Hypothyroid” means thyroid hormone synthesis is inhibited by 6-propyl-2-thiouracil (PTU), while “euthyroid” means the absence of inhibition. PTU was administered at a fixed concentration of 100 ppm. LT3, 3,3’,5-triiodo-L-thyronine; TRIAC, 3,3’,5-triiodothyroacetic acid.

To increase the sensitivity of our mouse model for determining TH actions, we induced hypothyroidism in mice using 6-propyl-2-thiouracil (PTU), an anti-thyroid drug that inhibits TH secretion (24). The hypothyroid mice we generated suffered growth retardation as age-dependent increases in body weight and naso-anal length were attenuated (Fig. 2A–D). This attenuation was not due to PTU toxicity but to weakened TH actions because growth retardation was rescued by co-administration of 0.1 ppm of either LT3 or TRIAC (Fig. 2A–D). Shortening of the lumbar spine and tibia was also restored by co-administration of either LT3 or TRIAC, which supports the involvement of PTU-induced growth retardation in skeletal growth (Fig. 2E–H).

**Fig. 2:**
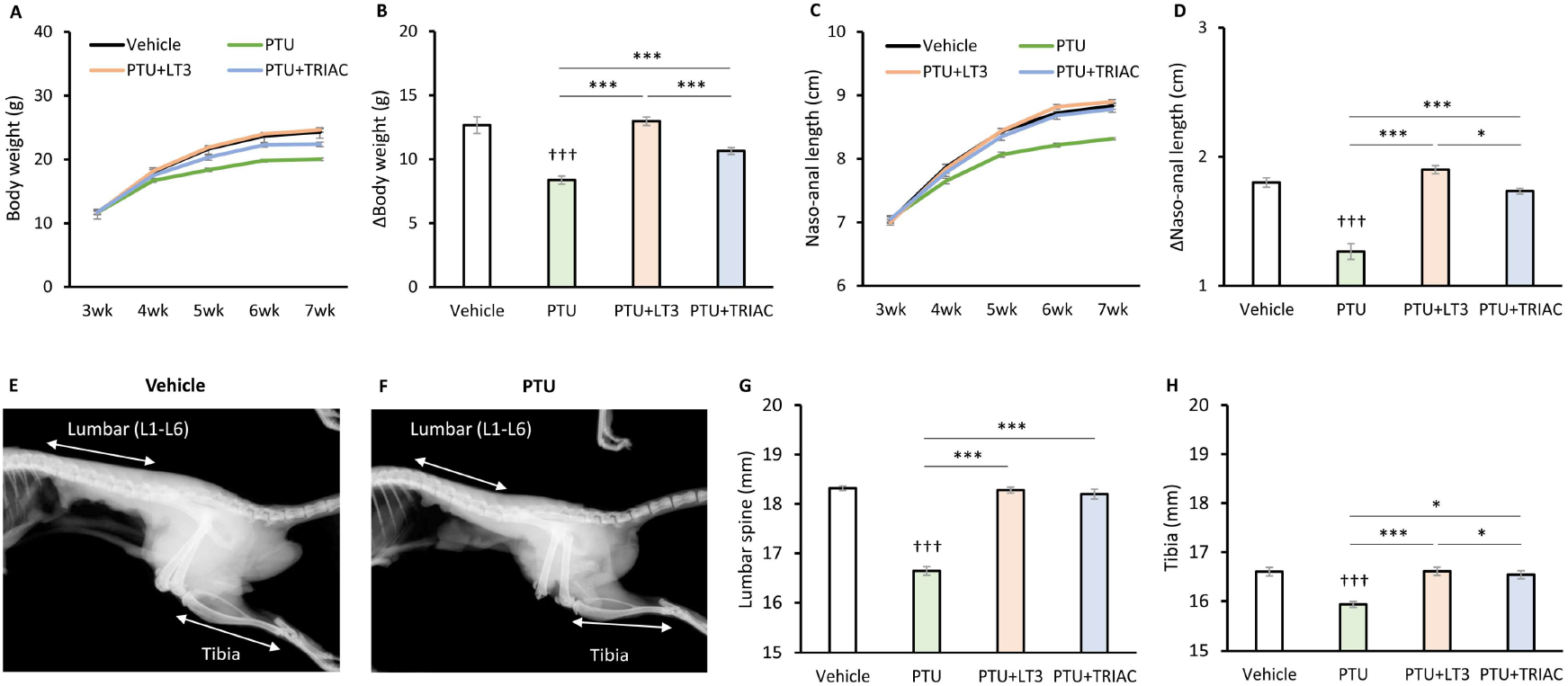
TRIAC effects on growth in hypothyroid mice. (A-D) Growth curves of hypothyroid mice co-administered with LT3 and TRIAC based on body weights (A) and naso-anal lengths (C), and relative amounts of change (Δ) (B, D). Vehicle, PTU, PTU + TRIAC, *n* = 6 each; and PTU + LT3, *n* = 5. (E, F) Soft X-ray images of mice administered with vehicle (E) and PTU (F) at seven weeks of age. (G, H) Bone lengths of hypothyroid mice at seven weeks of age measured on the soft X-ray images. Statistical analyses were performed using Student’s *t*-test for vehicle vs PTU and the *p*-value is presented as †p < 0.05, ††p < 0.01, and †††p < 0.001. One-way analysis of variance (ANOVA) followed by the Tukey-Kramer test was used for comparisons among PTU, PTU + LT3, and PTU + TRIAC experiments and the p-value is presented as \**p* < 0.05, \*\**p* < 0.01, and \*\*\**p* < 0.001.

The changes we observed in gross appearance suggest that hypothyroid mice are more suitable for determining TRIAC effects than euthyroid mice. We measured serum TH levels in hypothyroid mice and found that PTU administration decreased serum T4 and T3 levels, while LT3 co-administration increased serum T3 levels and TRIAC co-administration increased serum TRIAC levels (Fig. 3A–C). As for the HPT axis, PTU administration elevated serum TSH levels, which was feedback reaction to low serum T4 and T3 levels (Fig. 3D). Furthermore, we found that PTU administration enlarged the thyroid gland (Fig. 3E,F). Based on histological appearance, follicular cells thickened whereas follicle size decreased under PTU-induced TSH stimulation (Fig. 3G,H). These changes were fully abrogated by LT3 and reversed to a lesser extent by TRIAC co-administration (Fig. 3I,J).

**Fig. 3:**
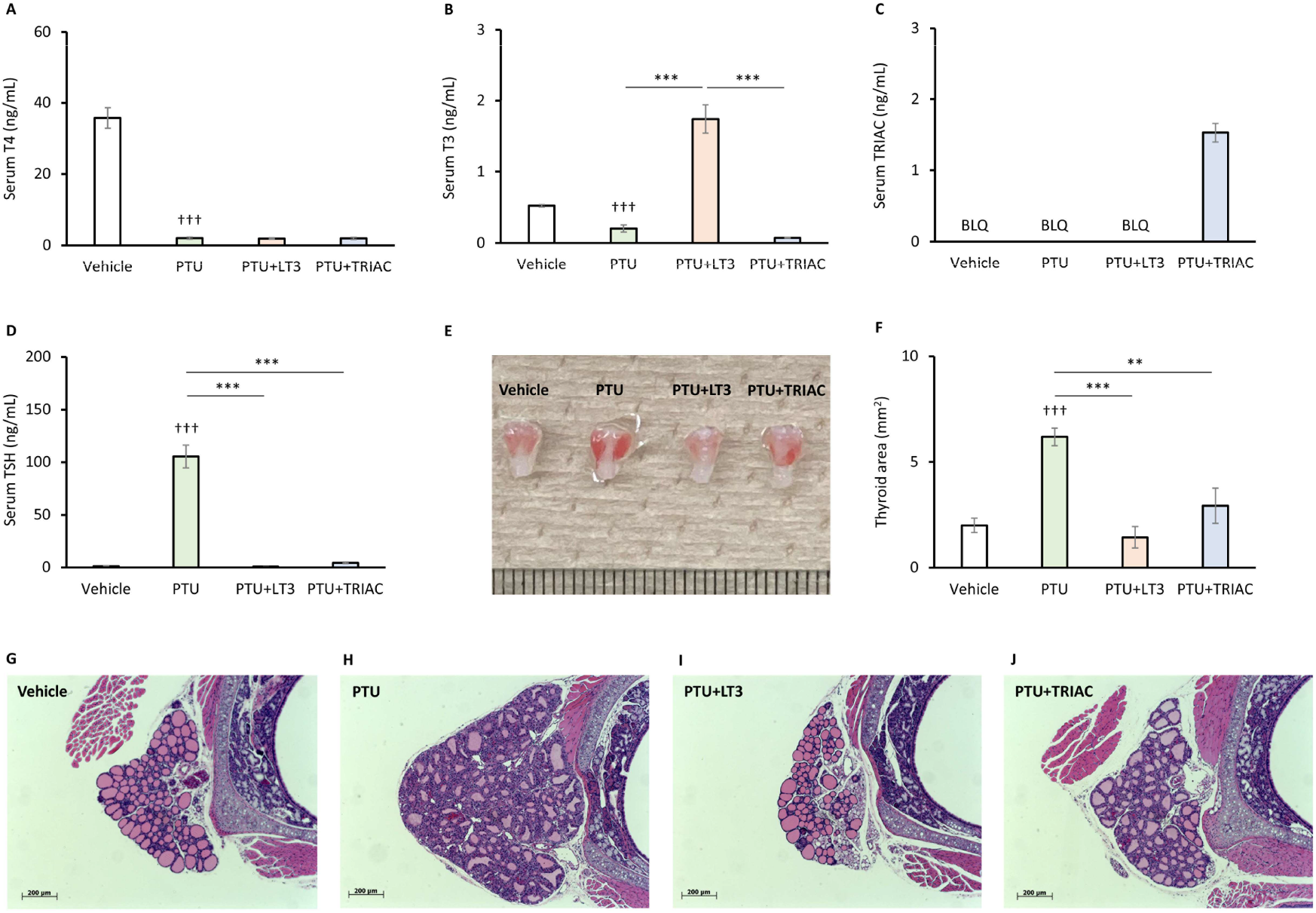
TRIAC effects on the hypothalamus-pituitary-thyroid axis in hypothyroid mice. (A-D) Serum levels of 3,3’,5,5’-tetraiodothyronine (T4), 3,3’,5-triiodothyronine (T3), TRIAC, and thyroid-stimulating hormone (TSH). (E) Gross appearance of thyroid glands. The scale on the ruler is 1 mm. (F) Thyroid gland size was measured as thyroid area on photograph. (G-J) Histological images of thyroid glands stained using hematoxylin and eosin. Vehicle, PTU, PTU + TRIAC, *n* = 6 each; PTU + LT3, *n* = 5. BLQ, below the limit of quantitation. Statistical analyses were performed as described in Fig. 2.

These results indicate that TRIAC can affect growth and the HPT axis in a similar manner to T3 even though TRIAC co-administration resulted in weaker changes compared with LT3 co-administration. To investigate differences between TRIAC and T3 with greater sensitivity, we measured the transcript levels of TH-responsive genes and regulators of TH action. As for the pituitary gland, an increase in *Tshb* (encoding TSH beta subunit) mRNA following PTU administration was abrogated by co-administering either LT3 or TRIAC (Fig. 4A), which corresponded to changes in serum TSH levels (Fig. 3D). PTU administration increased *Dio2* mRNA that upregulates TH action by generating T3 from 5’-deiodination of T4. On the other hand, PTU administration decreased *Dio3* mRNA that downregulates TH action by generating reverse T3 from 5-deiodination of T4 and 3,3’-diiodothyronine from 5-deiodination of T3 (Fig. 4A). Both changes were restored by co-administering either LT3 or TRIAC (Fig. 4A).

**Fig. 4:**
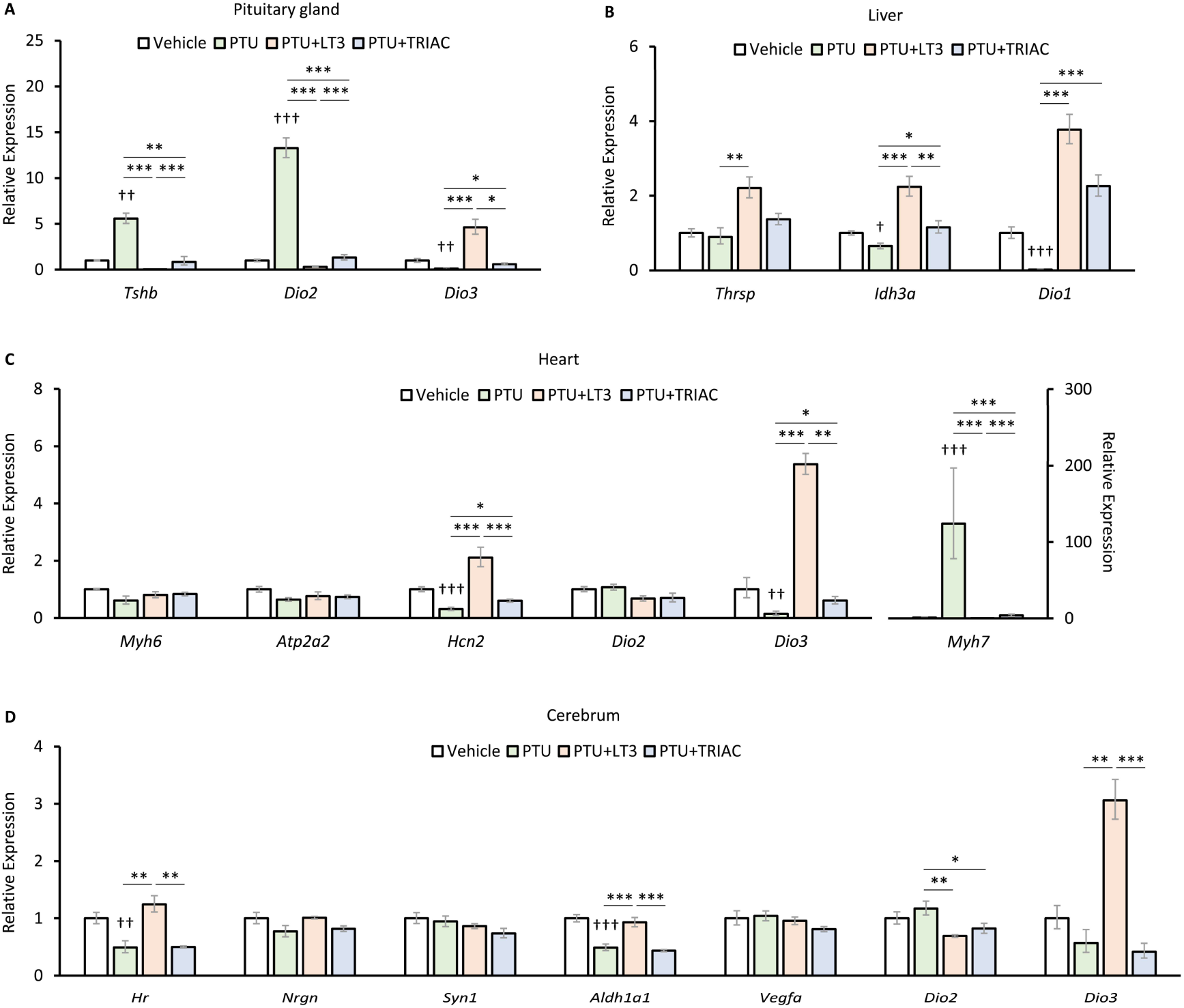
Gene expression profiles of hypothyroid mice determined by quantitative RT-PCR. (A) The pituitary gland, (B) the liver, (C) the heart, and (D) the cerebrum. Vehicle, PTU, PTU + TRIAC, *n* = 6 each; PTU + LT3, *n* = 5. Results are normalized using *Ppia* as an internal control and shown as fold change versus the vehicle-only control. Significance was confirmed if consistent results were observed using *Hprt* as another internal control (see Supplementary Fig. S2 for the results). Statistical analyses were performed as described in Fig. 2.

We subsequently measured mRNA levels in the liver, heart, and cerebrum. Liver mRNA levels of the TH-responsive genes *Thrsp, Idh3a,* and *Dio1* increased following LT3 co-administration and to a lesser extent, following TRIAC co-administration (Fig. 4B). In the heart, mRNA levels of the TH-responsive genes *Hcn2* and *Myh7* changed following PTU administration; *Hcn2* levels were lower while *Myh7* levels were higher (Fig. 4C). These changes were reversed by co-administering LT3 and to a lesser extent, following TRIAC co-administration. PTU administration also decreased *Dio3* mRNA levels in the heart (Fig. 4C). TRIAC co-administration reversed this change in *Dio3* mRNA abundance while LT3 co-administration increased *Dio3* mRNA abundance.

Unlike the liver and heart, we observed differences between TRIAC and T3’s effects on the cerebrum. Cerebral mRNA levels of the TH-responsive genes *Hr* and *Aldh1a1* were decreased following PTU administration; this decrease was reversed by LT3 co-administration but not by TRIAC co-administration (Fig. 4D). Additionally, *Dio3* mRNA levels were unchanged following TRIAC co-administration but were significantly higher following LT3 co-administration (Fig. 4D). We validated these results by normalizing them against an additional internal control, *Hprt* (Supplementary Fig. S2).

Based on our gene expression analysis, the cerebrum does not respond to TRIAC co-administration. To elucidate the underlying mechanisms, we measured TH levels in each organ. In the liver, PTU decreased T4, LT3 co-administration increased T3, and TRIAC co-administration increased TRIAC as expected (Fig. 5A–C). By contrast, in the cerebrum, TRIAC co-administration did not increase TRIAC levels, whereas LT3 co-administration increased T3 levels (Fig. 5D–F). In summary, TRIAC administration does not increase cerebral TRIAC abundance, and TH-responsive genes therefore do not respond.

**Fig. 5:**
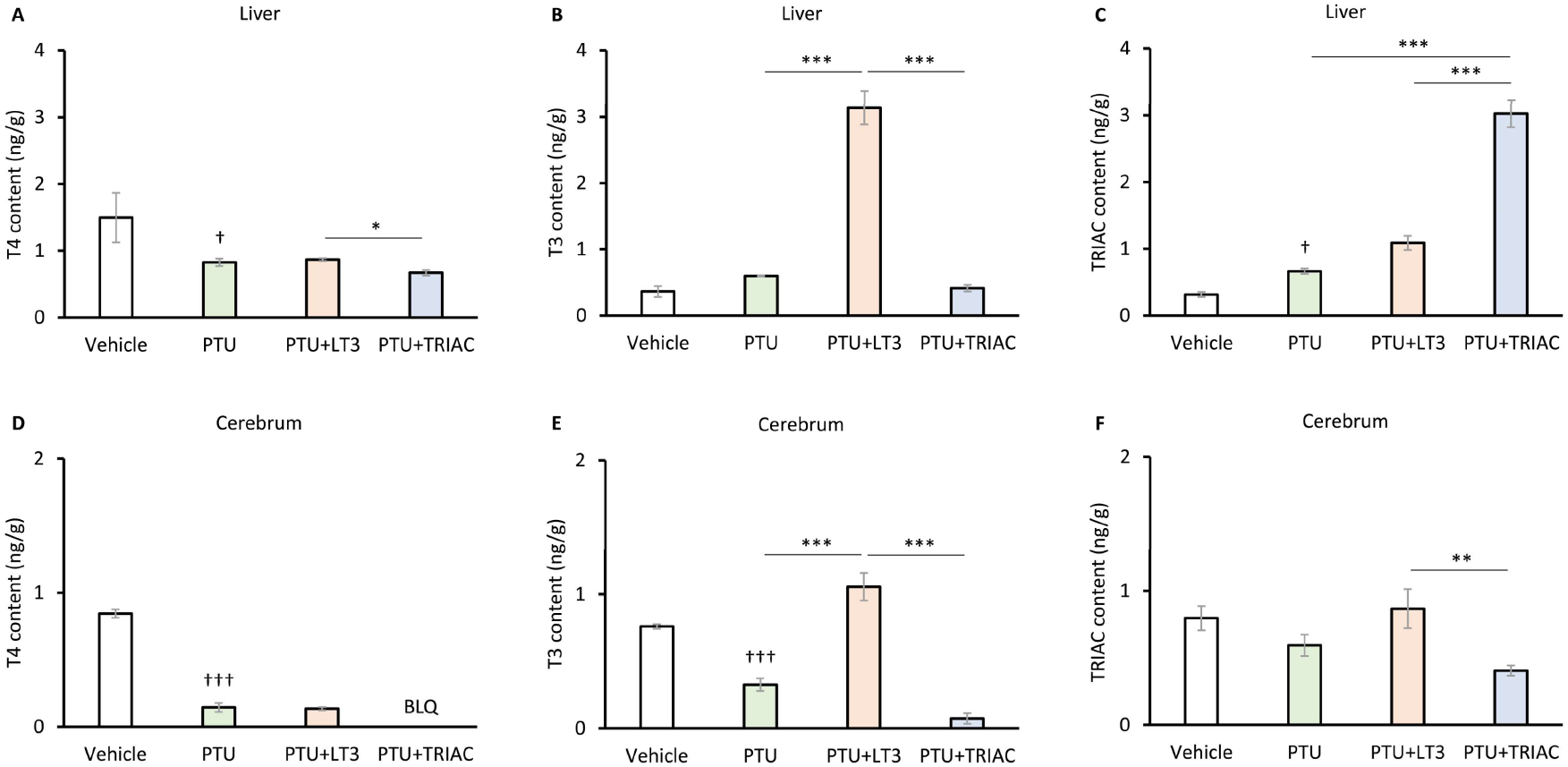
Organ-specific levels of thyroid hormones in hypothyroid mice. (A-C) Liver levels of T4 (A), T3 (B), and TRIAC (C). (D-F) Cerebral levels of T4 (D), T3 (E), and TRIAC (F). Results are presented as amount of each thyroid hormone per gram of organ weight. Vehicle, PTU, PTU + TRIAC, *n* = 6 each; PTU + LT3, *n* = 5. Statistical analyses were performed as described in Fig. 2.

### TRIAC administration to euthyroid mice

Based on the results of our experiments with hypothyroid mice, we hypothesized that exogenous TRIAC attenuates cerebral TH actions even in euthyroid mice. In detail, the supply of circulating T4 and T3 should be decreased by administering TRIAC, which suppresses the HPT axis (Fig. 3D–J). In addition, TRIAC does not appear to be transported into the cerebrum unlike T3 (Fig. 5E,F). We therefore predicted that TRIAC administration cannot rescue the attenuation of cerebral TH actions due to a reduction in cerebral T4 and T3.

To test our hypothesis, we further analyzed euthyroid mice with administering 0.1 ppm of either LT3 and TRIAC or 1 ppm (Fig. 1). We found that serum TH levels changed as expected in these mice (Fig. 6A–F). Serum T4 levels were decreased by 0.1 ppm LT3 and by both 1 ppm LT3 and 1 ppm TRIAC (Fig. 6A,D). Serum T3 levels were increased by 1 ppm LT3 but was decreased slightly by 0.1 ppm TRIAC and to a greater extent by 1 ppm TRIAC (Fig. 6B,E). Serum TRIAC levels were increased by TRIAC administration in a dose-dependent manner (Fig. 6F) and become slightly detectable by LT3 at 1 ppm (Fig. 6C). We adopted 1 ppm as a working concentration because we verified substantial reduction in serum T4 and T3 levels following the administration of 1 ppm TRIAC.

**Fig. 6:**
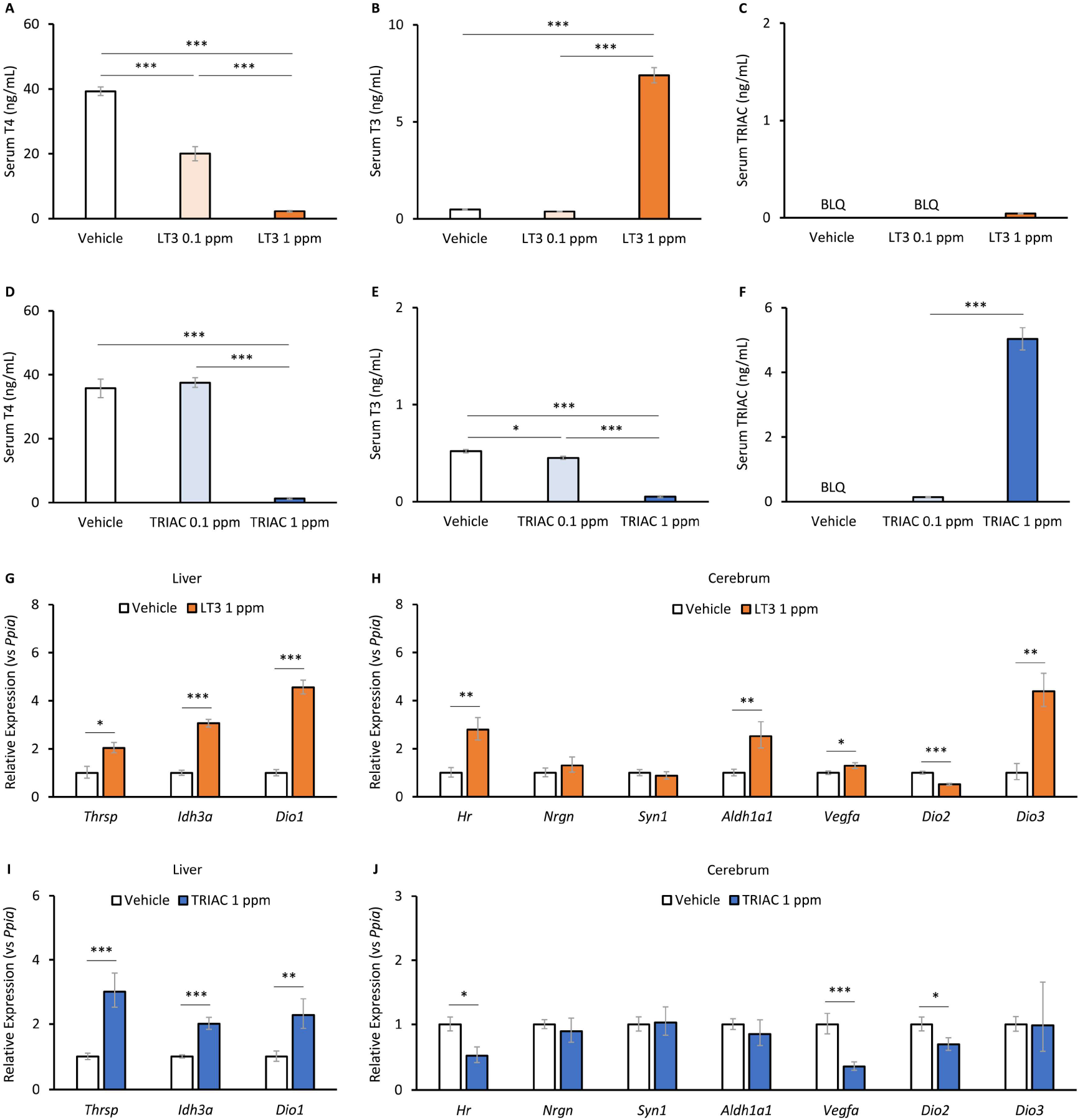
Serum thyroid hormone levels and gene expression profiles of euthyroid mice. (A-C) Serum levels of T4, T3, and TRIAC in mice administered with LT3 (n = 6 each). (D-F) Serum levels of T4, T3, and TRIAC in mice administered with TRIAC (n = 6 each). (G-J) Gene expression profiles determined by quantitative RT-PCR; liver (G) and cerebrum (H) of mice administered with LT3 at 1 ppm (n = 6 each); liver (I) and cerebrum (J) of mice administered with TRIAC at 1 ppm (*n* = 6 each). Results are normalized using *Ppia* as an internal control and shown as fold change versus the vehicle-only control. Significance was verified if consistent results were observed using *Hprt* as an alternative internal control (see Supplementary Fig. S3 for the results). Statistical analyses were performed by Student’s t-test for panels G-J and by ANOVA followed by the Tukey-Kramer test for panels A-F. \**p* < 0.05, \*\**p* < 0.01, and \*\*\**p* < 0.001.

We subsequently measured gene expression in the liver and cerebrum. LT3 administration increased the abundance of TH-responsive gene transcripts in both the liver and cerebrum (Fig. 6G,H). Similarly to hypothyroid mice (Fig. 4D), the abundance of *Dio2* mRNA in the cerebrum decreased while that of *Dio3* mRNA increased following 1 ppm LT3 administration (Fig. 6H). Meanwhile, TRIAC administration did not increase the abundance of TH-responsive gene transcripts in the cerebrum. Instead, *Hr* and *Vegfa* mRNA levels were lower even though the mRNAs of TH-responsive genes were significantly higher in the liver (Fig. 6I,J). Furthermore, cerebral *Dio3* mRNA levels were unchanged following TRIAC administration (Fig. 6J). We validated these results by normalizing against *Hprt* (Supplementary Fig. S3).

We proceeded to measure organ-specific TH levels to determine whether quantitative changes in local THs may be responsible for the observed downregulation of cerebral TH-responsive genes in response to TRIAC. Similarly to serum levels (Fig. 6A–F), liver T3 and T4 levels were both altered by LT3 and by TRIAC (Fig. 7A–F). Liver T4 levels were decreased by both 1 ppm LT3 and 1 ppm TRIAC (Fig. 7A,D). Liver T3 abundance was increased by 1 ppm LT3 but unchanged by 1 ppm TRIAC (Fig. 7B,E). Liver TRIAC abundance was increased by TRIAC administration in a dose-dependent manner but unchanged by LT3 (Fig. 7C,F). We also detected changes in cerebral T3 and T4 levels. LT3 administration decreased cerebral T4 abundance, increased T3 abundance, and decreased TRIAC abundance (Fig. 7G–I). By contrast, cerebral T4 and T3 levels fell following TRIAC administration while TRIAC levels remained unchanged (Fig. 7J–L).

**Fig. 7:**
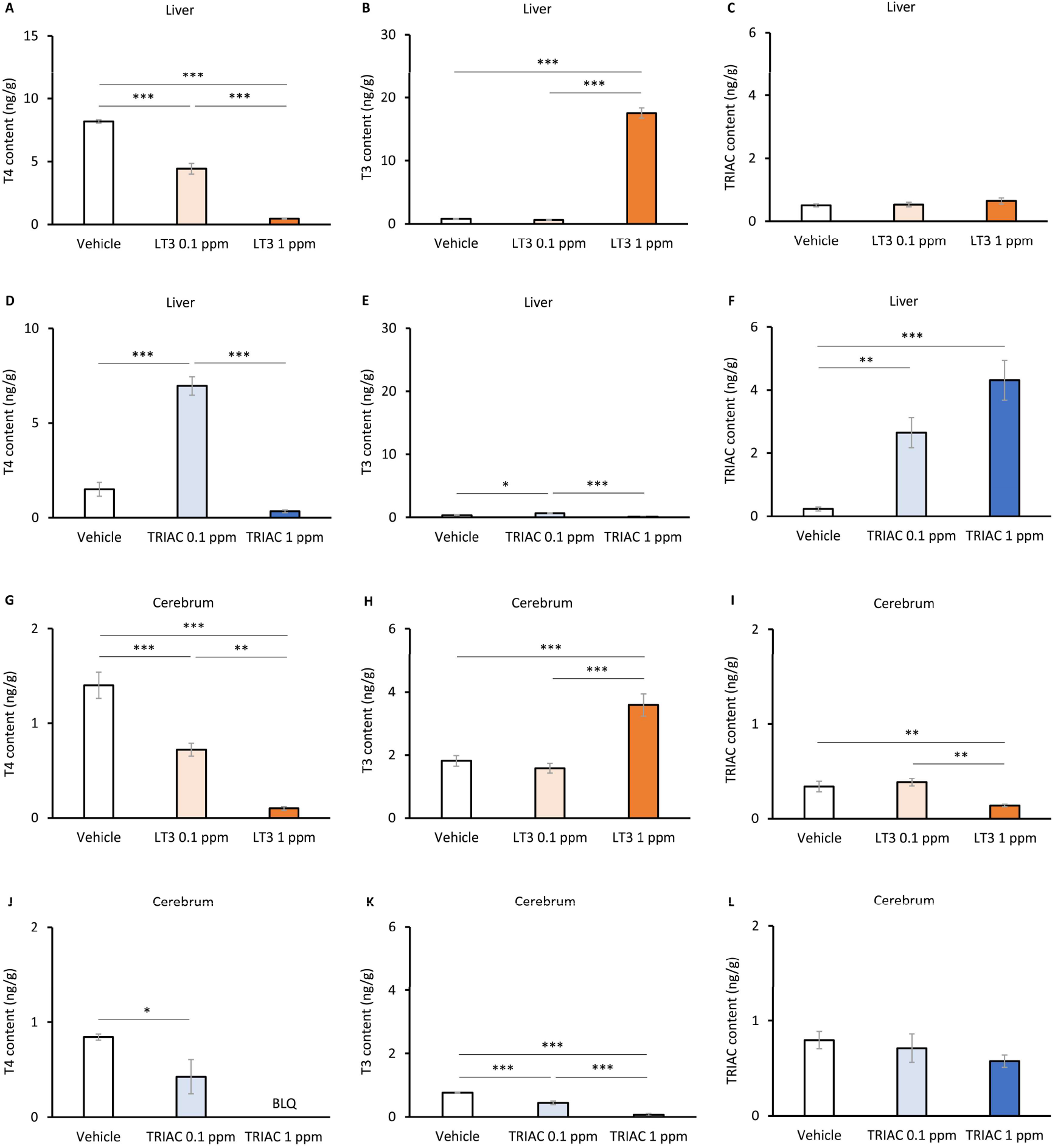
Organ-specific levels of thyroid hormones in euthyroid mice. (A-F) Liver-specific levels of T4 (A), T3 (B), and TRIAC (C) in mice administered with LT3 (*n* = 6 each) and those of T4 (D), T3 (E), and TRIAC (F) in mice administered with TRIAC (*n* = 6 each). (G-L) Cerebral levels of T4 (G), T3 (H), and TRIAC (I) in mice administered with LT3 (*n* = 6 each) and those of T4 (J), T3 (K), and TRIAC (L) in mice administered with TRIAC (*n* = 6 each). Results are presented as amount of each thyroid hormone per gram of tissue weight. Statistical analyses all involved ANOVA followed by the Tukey-Kramer test. \**p* < 0.05, \*\**p* < 0.01, and \*\*\**p* < 0.001.

We further examined elements that regulate TH actions to verify that the HPT axis mediates changes in overall and organ-specific levels of THs following TRIAC administration. We observed significant changes in liver mRNA levels of *Thra* and *Thrb,* which encode THRs, as well as *Mct8,* a TH transporter. The levels of these three transcripts were higher following LT3 administration but lower after TRIAC administration (Supplementary Fig. S4A–D). Nevertheless, TRIAC upregulated liver TH-responsive genes (Fig. 6I). As for genes that mediate the glucuronidation of THs, *Ugt1a1* mRNA was increased by LT3 but not significantly changed by TRIAC, whereas *Ugt1a9* mRNA was decreased by both LT3 and TRIAC (Supplementary Fig. S4A–D). The serum TH-lowering effects of TRIAC are apparently not due to glucuronidation in the liver. The only significant change observed in the cerebrum was a decrease in *Thra* mRNA following LT3 administration (Supplementary Fig. S4E–H). The mRNA levels of *Oatp1c1*, another TH transporter, and *Mct8* did not change in response to either LT3 or TRIAC. As for the pituitary gland, mRNA levels of *Tshb* and *Dio2* were decreased and *Dio3* mRNA was increased by administering either LT3 or TRIAC (Supplementary Fig. S5A). We validated these results by normalizing them against an additional internal control, *Hprt* (Supplementary Fig. S5B). Moreover, administration of either LT3 or TRIAC reduced thyroid gland size (Supplementary Fig. S5C,D) and thickness of follicular cells (Supplementary Fig. S5E–G).

## Discussion

As environmental water could be contaminated with TRIAC (18), we determined the biological effects of exogenous TRIAC. Comparisons were performed against T3, a main endogenous contributor of TH actions. By administering either TRIAC or T3 to hypothyroid mice, we found both compounds exerted similar effects on growth and the HPT axis. Specifically, TRIAC upregulated TH-responsive genes of the pituitary gland, the liver, and the heart, as did T3. Interestingly, we observed that TRIAC could not rescue the downregulation of TH-responsive genes in the cerebrums of hypothyroid mice. By measuring organ-specific TH levels, we found that TRIAC was not sufficiently distributed into the cerebrum when administered orally. Altogether, we verified that TRIAC administration attenuates cerebral TH actions even in the euthyroid state. The mechanism underlying TH depletion in the cerebrum appears to involve suppression of the HPT axis and disabling TRIAC delivery to the cerebrum.

Euthyroid models have conventionally been used to evaluate EDCs (1, 25). In this study, we induced hypothyroidism in mice using PTU to investigate effects on TH action with high sensitivity. Growth effects could not be determined in euthyroid mice but could be determined in hypothyroid mice as growth retardation was restored by LT3 and TRIAC administration. Moreover, hypothyroid mice visualized changes in the HPT axis as enlargement of the thyroid gland and follicular dysmorphology. This observation reflects the stimulating effects of elevated circulating TSH via feedback reaction against TH depletion. In addition, hypothyroid mice enabled changes in TH-responsive gene expression to be measured clearly. We believe that other EDCs and candidates can be further elucidated using hypothyroid mice.

We could interpret TH-responsive gene profiles using the results of controls, namely PTU administration for downregulation and LT3 co-administration for upregulation. We verified the upregulatory effects of TRIAC on multiple TH-responsive genes in the pituitary gland, liver, and heart. In the cerebrum, *Hr* and *Aldh1a1* mRNA levels were decreased by PTU administration and this decrease was rescued by LT3 co-administration. However, TRIAC co-administration could not restore the downregulation of *Hr* and *Aldh1a1*. We observed a similar discrepancy for *Dio3;* both LT3 and TRIAC co-administration upregulated *Dio3* mRNA levels in the heart, whereas only LT3 upregulated *Dio3* in the cerebrum. DIO3 decreases cellular T3 by 5’-deiodination and its activity in the cerebral cortex is increased in hyperthyroid rats and decreased in hypothyroid rats (26).

Gene expression profiles of hypothyroid mice indicated that exogenous TRIAC does not augment TH actions in the cerebrum. The underlying mechanism was elucidated by comparing the TH levels of the liver and cerebrum. Liver T3 and TRIAC levels increased following administration of the respective compound. By contrast, cerebral TRIAC levels were not increased by its administration although cerebral T3 levels were increased by LT3 administration. Thus, TRIAC does not appear to be trafficked into the cerebrum, which could explain why cerebral TH-responsive genes are unresponsive to exogenous TRIAC.

T4 and T3, major endogenous THR agonists, cross plasma membranes via transporters such as the monocarboxylate transporters (MCTs), organic anion– transporting polypeptides (OATPs), L-type amino acid transporters, and sodium/taurocholate cotransporting polypeptide (27). MCT8 is an essential transporter of T4 and T3 particularly in central nervous system (CNS). Patients with mutations in *MCT8* develop Allan–Herndon–Dudley syndrome, which is characterized by a severe neurodevelopmental defect as well as thyroid dysfunction (28). *Mct8*-knockout mice have less T4 and T3 in their brains compared with wild type (29). Although OATP1C1 concentration within the human blood–brain barrier (BBB) is much lower than in the murine BBB (30), OATP1C1 can transport THs into the brain because brain levels of T4 and T3 are even lower in *Mct8/Oatp1c1* double knockout mice compared with *Mct8* knockout mice (31).

The potential to be transported by either MCT8, OATP1C1, or both is quite important for TH action in CNS of EDCs because MCT8 and OATP1C1 are both expressed in the human BBB (30). Importantly, TRIAC is apparently not transported by MCT8 (32). Although there have been no reports on the relationship between TRIAC and OATP1C1, TETRAC, a precursor hormone of TRIAC, is not transported by either MCT8 or OATP1C1 (33). In this context, insufficient CNS distribution of exogenous TRIAC was assumed to be due to poor permeability through the BBB. This assumption was supported not only by our results but also by other reports that TRIAC administration does not upregulate TH-responsive genes in the mouse brain (34, 35).

The results from hypothyroid mice raised the concern that exogenous intake of TRIAC can disrupt TH actions in the CNS as demonstrated in Fig. 8. First, exogenous TRIAC decreases TH secretion from the thyroid gland by suppressing the HPT axis. Second, the decrease in circulating TH levels caused by TRIAC leads to insufficient TH levels and attenuation of TH actions in various tissues. Third, exogenous TRIAC compensates for TH actions by acting as an alternative to endogenous THs in peripheral tissues. However, TRIAC does not compensate for TH actions in the CNS because TRIAC hardly distributes through the BBB. We tested our hypothesis by analyzing euthyroid mice administered TRIAC. TRIAC administration decreased serum and liver levels of T4 and T3 but transcript levels of TH-responsive genes were increased in the liver. Meanwhile, TRIAC administration failed to increase TRIAC levels and instead, downregulated TH-responsive genes in the cerebrum. In other words, TRIAC administration depleted T4 and T3 from the cerebrum without TRIAC being trafficked into the cerebrum, which resulted in the downregulation of TH-responsive genes in the cerebrum. Additionally, we determined that the HPT axis is the main contributor to the depletion of THs by TRIAC administration and that the expression of genes encoding THRs and transporters did not change significantly in the cerebrum.

**Fig. 8:**
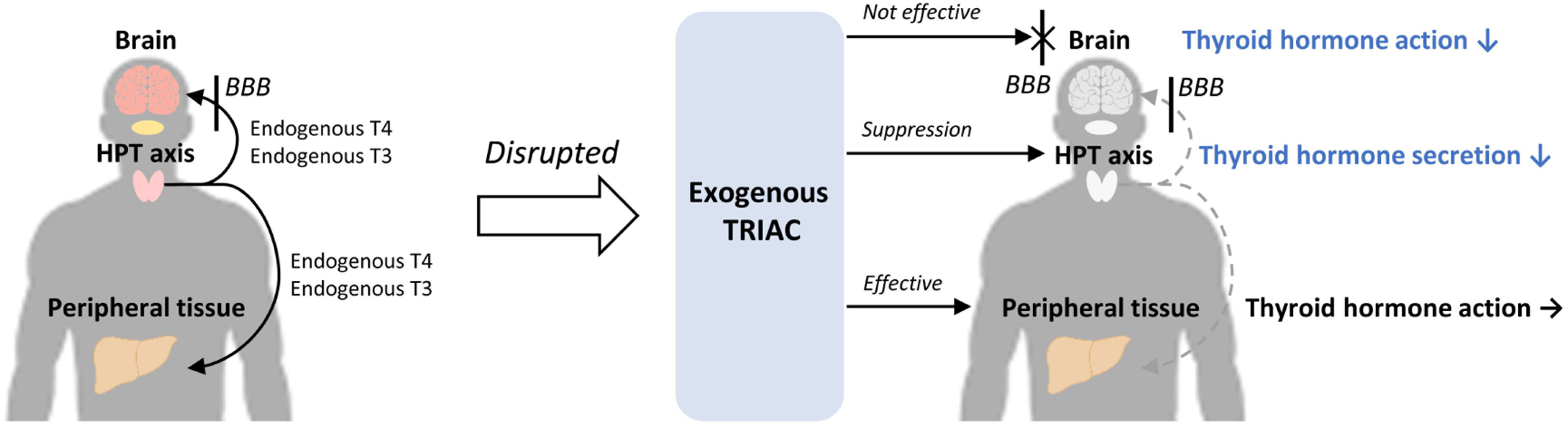
Schematic summary for putative disruption mechanisms of TRIAC. BBB, blood–brain barrier.

The mechanisms of EDC action have been proposed in an expert consensus statement (8). Of the characteristics presented in the statement, our observations suggest that TRIAC exerts its disruptive effects by activating hormone receptors, interfering with hormone synthesis, and altering circulating hormone levels. We also determined that these multiple characteristics contribute cooperatively to the disruptive effects of TRIAC. Furthermore, the key disruptive characteristic of TRIAC is its heterogenous distribution among organs, specifically its poor permeability across the BBB. We have therefore identified a novel concept of EDC action.

As described in the Introduction, the effluent from several sewage treatment plants have been found to contain TRIAC, which suggests that environmental water can be contaminated by TRIAC (18). In wildlife, malformations were developed not only by T4 and T3 but by exposure to TRIAC (36). The data we have presented here provides a warning of an adverse risk posed by environmental TRIAC towards the human CNS. THs play crucial roles in fetal and neonatal neurodevelopment because profound and permanent CNS defects have been observed in patients with untreated congenital hypothyroidism, resistance to thyroid hormone α (37, 38), and Allan–Herndon–Dudley syndrome (28). Even milder attenuation of TH actions in the CNS could pose a significant risk to neurodevelopment. For example, children with permanent congenital hypothyroidism were reported to have lower cognitive scores than those with transient congenital hypothyroidism and control subjects even if they received replacement therapy (39).

The measurement of TRIAC in various settings is important for further investigation. TRIAC levels in umbilical cord blood may directly suggest a neurodevelopment risk. Similarly, serum TRIAC levels or the ratio of TRIAC to other THs may be a biomarker for cognitive risk both in children and adults. The impact of long-term low-dose exposure can be evaluated by an ongoing birth cohort study with environmental TRIAC monitoring.

In conclusion, exogenous TRIAC intake can attenuate TH actions exclusively in the cerebrum. The disruptive mechanisms of TRIAC were characterized as the additive effect of circulating endogenous THs being depleted via negative feedback acting on the HPT axis, THR activation in peripheral tissues, and poor permeability across the BBB. Our finding gives caution for environmental TRIAC and offers epidemiological studies in combination with TRIAC measurement.

## Materials and Methods

### Animals and chemicals

Three-week-old C57BL/6J male mice were purchased from Japan SLC, Inc. (Hamamatsu, Japan). PTU (Tokyo Chemical Industry Co., Ltd., Tokyo Japan), LT3 sodium salt (Sigma-Aldrich, St. Louis, MO), and TRIAC (Sigma-Aldrich) were purchased for administration to mice. All experiments involving mice were approved by the Animal Research Committee, Graduate School of Medicine, Kyoto University (permit number: Med Kyo 21239). Animal care and experiments were conducted in accordance with our institutional guidelines.

### Chemical administration to mice and sample collection

Mice were administered the chemicals via drinking water from 3 to 7 weeks of age. Concentrations of PTU, LT3, and TRIAC used were 100 ppm, 0.1 ppm or 1 ppm, and 0.1 ppm or 1 ppm, respectively. All mice were sacrificed by isoflurane exposure. To obtain sera, blood samples were immediately collected from the inferior vena cava, left undisturbed for 45 min at room temperature, and subsequently cooled on ice. After centrifugation at 3,000 *g* for 15 min at 4°C, the supernatants were collected and stored at −80°C until use. Thyroid glands harvested for histological analyses were first fixed in 4% paraformaldehyde phosphate buffer solution (FUJIFILM Wako Pure Chemical Corporation, Osaka, Japan), embedded in paraffin, and stained using hematoxylin and eosin. Thyroid gland size was measured as thyroid area on photograph using ImageJ software (National Institutes of Health, Bethesda, MD). Lengths of tibia and lumbar vertebrae were measured on soft X-ray film. Pituitary glands and thyroid glands harvested for RNA extraction were preserved in RNAprotect Tissue Reagent (QIAGEN, Venlo, Netherlands) at −20°C until use. Livers, hearts, and cerebrums were collected in microtubes, immediately frozen in liquid nitrogen, and stored at −80°C until use.

### Thyroid hormone measurement of serum

We measured serum levels of T4, T3, and TRIAC using liquid chromatography-tandem mass spectrometry (LC-MS/MS). Twenty microliters of each serum sample was mixed with 10 μL internal standard (IS) solution and 100 μL acetonitrile using a vortex mixer. The mixture was centrifuged at 15,000 rpm for 10 min at 4°C. T4-^13^C_6_ and T3-^13^C_6_ were used at 10 ng/mL as ISs for T4 and T3 measurements, respectively. For TRIAC measurements, T3-^13^C_6_ was used as the IS. Injection samples consisted of 60 μL of the supernatants and 240 μL of water. We injected 20 μL of each prepared sample into a Nexera X2 UHPLC system (Shimadzu Corporation, Kyoto, Japan) with a Raptor Biphenyl column (2.1 × 50 mm, 1.8 μm; Restek, Bellafonte, PA) maintained at 40°C. A gradient of mobile phase A (0.1% FA in water) and mobile phase B (methanol) was used. General conditions were as follows: 60% methanol (0–2.5 min), 60–80% methanol linear gradient (2.5–2.6 min), 80% linear gradient (2.6–4 min), 95% linear gradient (4–6 min), 60% linear gradient (6–8 min); flow rate of 0.4 mL/min (0–4 min), 0.5 mL/min (4–6 min), 0.4 mL/min (6–8 min). The following MS settings were adopted: curtain and collision gas pressure of 32 and 9 psi, respectively; ion spray voltage of 2500 V (positive mode) and −2000 V (negative mode); temperature of 450°C; and ion source gas 1 and 2 pressure of 40 and 70 psi; correspondingly. A Triple Quad™ 7500 system (AB SCIEX, Tokyo, Japan) was used to measure T4 and T3 with electrospray ionization (ESI) in positive mode and TRIAC with ESI in negative mode by multiple reaction monitoring (MRM).

Serum TSH levels were evaluated using a Rodent TSH ELISA TEST KIT (Endocrine technologies, Inc., Newark, CA) and 20 μL of serum from each mouse. Absorbance was determined using an iMark microplate reader (Bio-Rad, Hercules, CA).

### Thyroid hormone measurement of organ

Organ-specific T4, T3, and TRIAC levels were measured using LC-MS/MS. We mechanically homogenized 50 mg of either the liver or cerebrum of a mouse in 300 μL radioimmunoprecipitation buffer (Nacalai Tesque, Kyoto, Japan) and left the homogenate on ice for 30 min. Supernatants were centrifuged at 10,000 *g* for 10 min at 4°C and collected in microtubes. We added 300 μL methanol and 600 μL chloroform, mixed with vortex mixer, and centrifuged at 15,000 *g* for 2 min at 4°C. The upper water/methanol phase was collected in new microtubes. We injected 6 μL of each sample into a Triple Quad™ 5500+ system and QTRAP Ready (AB SCIEX). Chromatography was performed using InertSustain C18 column (2.1 × 150 mm, 5 μm;

GL Sciences, Tokyo, Japan) maintained at 40°C. A gradient of mobile phase A (0.5 mM ammonium fluoride in water) and mobile phase B (methanol) was used. The general conditions were as follows: 40% methanol (0–1 min), 40–90% methanol linear gradient (1–10 min), 90% linear gradient (10–15 min); flow rate of 0.2 mL/min. The following MS settings were adopted: curtain and collision gas pressure of 40 and 8 psi, respectively; ion spray voltage of −4500 V; temperature of 500°C; and ion source gas 1 and 2 pressure of 80 and 70 psi; correspondingly. T4, T3, and TRIAC were all measured with ESI in negative mode by MRM.

### RNA extraction and quantitative RT-PCR

Total RNA was extracted using a Nucleospin RNA Plus kit (Macherey-Nagel, Düren, Germany) and reverse transcribed using ReverTra Ace (TOYOBO Life Science, Osaka, Japan). For cerebrums, we extracted total RNA using QIAzol Lysis Reagent (QIAGEN) and subsequently cleaned up with a Nucleospin RNA Plus kit. Quantitative RT-PCR was performed using THUNDERBIRD SYBR qPCR MIX (TOYOBO Life Science) with the StepOnePlus Real-Time PCR System (Thermo Fisher Scientific, Waltham, MA). Results were normalized using *Ppia* and *Hprt* as the reference gene; relative mRNA expression of target genes was evaluated using the comparative threshold cycle method. Primer sequences used are listed in Supplementary Table S1.

### Statistical analysis

Results are expressed as means ± standard error of the mean and statistical analysis was performed using either Student’s t-test or one-way analysis of variance followed by the Tukey-Kramer test. JMP Pro version 16.1.0 (SAS Institute Inc., Cary, NC) was used for all statistical analyses. Statistical significance was defined as a *p-*value < 0.05.

## Acknowledgements

This work was supported by the Environment Research and Technology Development Fund (JPMEERF20195053) of the Environmental Restoration and Conservation Agency of Japan.

