## Supplementary Table S1 and Fig. S1-S5 for "TRIAC disrupts cerebral thyroid hormone action via a negative feedback loop and heterogenous distribution among organs"

**Supplementary Table S1. Primers used for quantitative RT-PCR**

| Genes | Primers |
|---|---|
| *Tshb* (NM_009432) | Fw: GAGAGTGTGCCTACTGCCTG<br>Rv: ACAGACATCCTGAGAGAGTGC |
| *Dio2* (NM_010050) | Fw: CTTCCTCCTAGATGCCTACAAAC<br>Rv: CGAGGCATAATTGTTACCTGATTC |
| *Dio3* (NM_172119) | Fw: CCGCTCTCTGCTGCTTCAC<br>Rv: CGGATGCACAAGAAATCTAAAAGC |
| *Thrsp* (NM_009381) | Fw: TCGGGGTCTTCATCAGTCTT<br>Rv: GCGGAAATACCAGGAAATGA |
| *Idh3a* (NM_029573) | Fw: ACGAGATGTACCTTGATACTG<br>Rv: ACACAGATCACTAAGGATGTC |
| *Dio1* (NM_007860) | Fw: CATCTGGGATTTCATTCAAGGC<br>Rv: TGGAGGCAAAGTCATCTACGAGTC |
| *Myh6* (NM_010856) | Fw: CAGACAGAGATTTCTCCAACCCA<br>Rv: GCCTCTAGGCGTTCCTTCTC |
| *Atp2a2* (NM_009722) | Fw: AACTACCTGGAACAACCCGC<br>Rv: TCATGCAGAGGGCTGGTAGA |
| *Hcn2* (NM_008226) | Fw: CCAGTCCCTGGATTCGTCAC<br>Rv: TCACAATCTCCTCACGCAGT |
| *Myh7* (NM_080728) | Fw: CACGTTTGAGAATCCAAGGCTC<br>Rv: CTCCTTCTCAGACTTCCGCA |
| *Hr* (NM_021877) | Fw: AAGCTAAATAGGGGATCCTG<br>Rv: ATTTGTAGAACGGACCACAC |
| *Nrgn* (NM_022029) | Fw: GCCAGACGACGATATTCTTGAC<br>Rv: TATCTTCTTCCTCGCCATGTGG |
| *Syn1* (NM_013680) | Fw: TGTGCGTGTCCAGAAGATTG<br>Rv: ACATGGCAATCTGCTCAAGC |
| *Aldh1a1* (NM_013467) | Fw: AAAATGTCTCCATCACTTGG<br>Rv: AAGTCTTTGCCAATGCATAC |
| *Vegfa* (NM_009505) | Fw: AACGAACGTACTTGCAGATG<br>Rv: GTGACATGGTTAATCGGTC |
| *Thra* (NM_178060) | Fw: CATGGACTTGGTTCTAGATG<br>Rv: CTGTAGCAACATGTATCAGG |
| *Thrb* (NM_009380) | Fw: GAGACTCTAACTTTGAATGGG<br>Rv: CGATCTGAAGACATTAGCAG |
| *Mct8/Slc16a2* (NM_009197) | Fw: CGTGCACCTGATGAAATATG<br>Rv: GATCATCATGGACATCAAGC |

| | |
|---|---|
| *Ugt1a1* (NM_201645) | Fw: TCTGAGCCCTGCATCTATCTG<br>Rv: CCCCAGAGGCGTTGACATA |
| *Ugt1a9* (NM_201644) | Fw: GAAGAACATGCATTTTGCTCCT<br>Rv: CTGGGCTAAAGAGGTCTGTCATAGTC |
| *Oatp1c1/Slco1c1* (NM_021471) | Fw: GGGCCATCCTTTACAGTCGG<br>Rv: CCTTCTCTCTATCTGAGTCACGG |
| *Ppia* (NM_008907) | Fw: CGCGTCTCCTTCGAGCTGTTTG<br>Rv: TGTAAAGTCACCACCCTGGCACAT |
| *Hprt* (NM_013556) | Fw: GGACCTCTCGAAGTGTTGGATAC<br>Rv: GCTCATCTTAGGCTTTGTATTTGGCT |

Fw: forward primer (5′–3′), Rv: reverse primer (5′–3′)

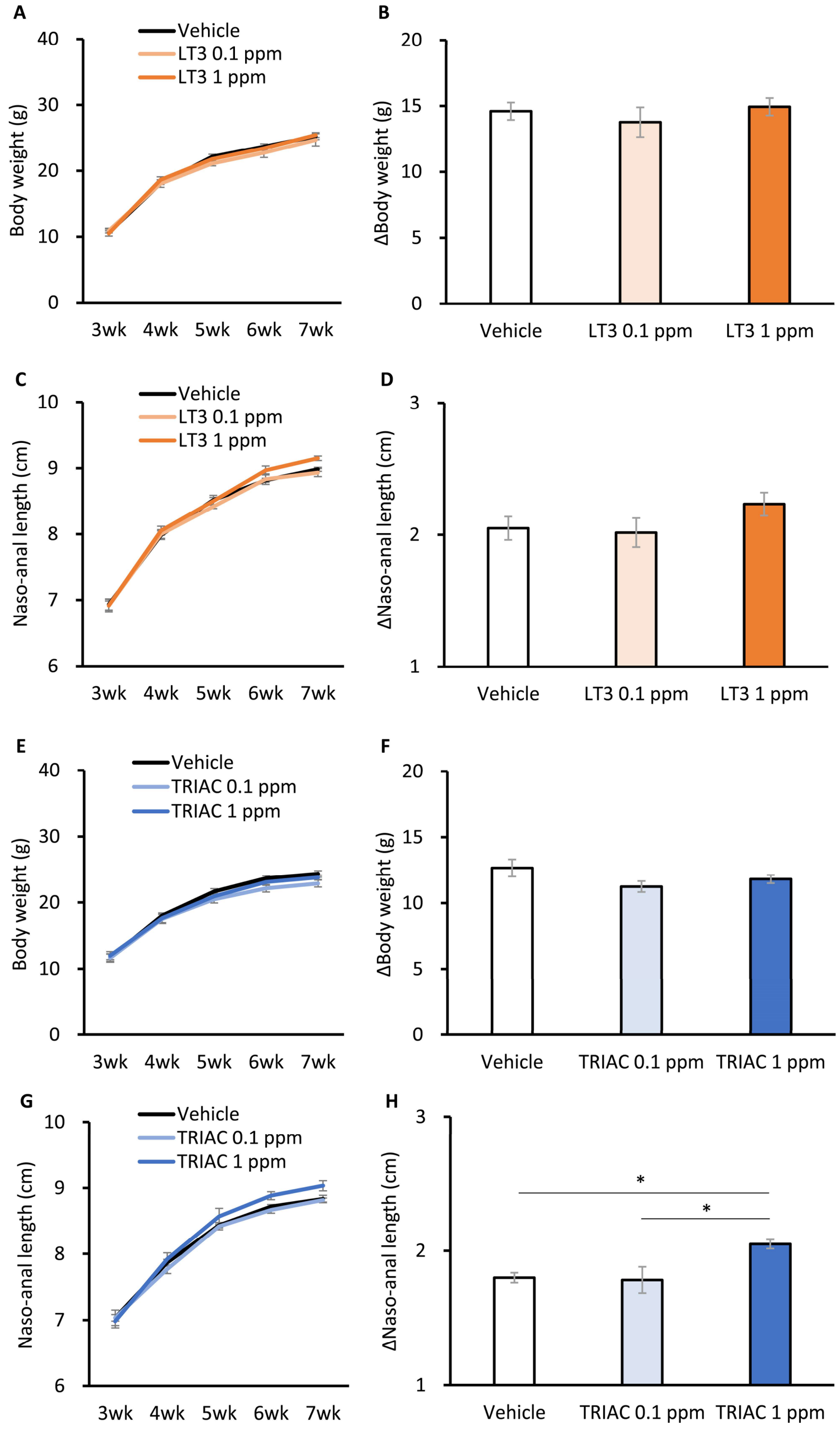


**Supplementary Fig. S1:** (A-D) Growth curves of euthyroid mice administered with 3,3′,5-triiodo-L-thyronine (LT3) based on body weights (A) and naso-anal lengths (C), and relative amounts of change (Δ) (B, D) (*n* = 6 each). (E-H) Growth curves of euthyroid mice administered with 3,3′,5-triiodothyroacetic acid (TRIAC) based on body weights (E) and naso-anal lengths (G), and their Δ (F, H) (*n* = 6 each). One-way analysis of variance (ANOVA) followed by the Tukey-Kramer test was used for comparisons and the p-value is presented as *$p < 0.05$.

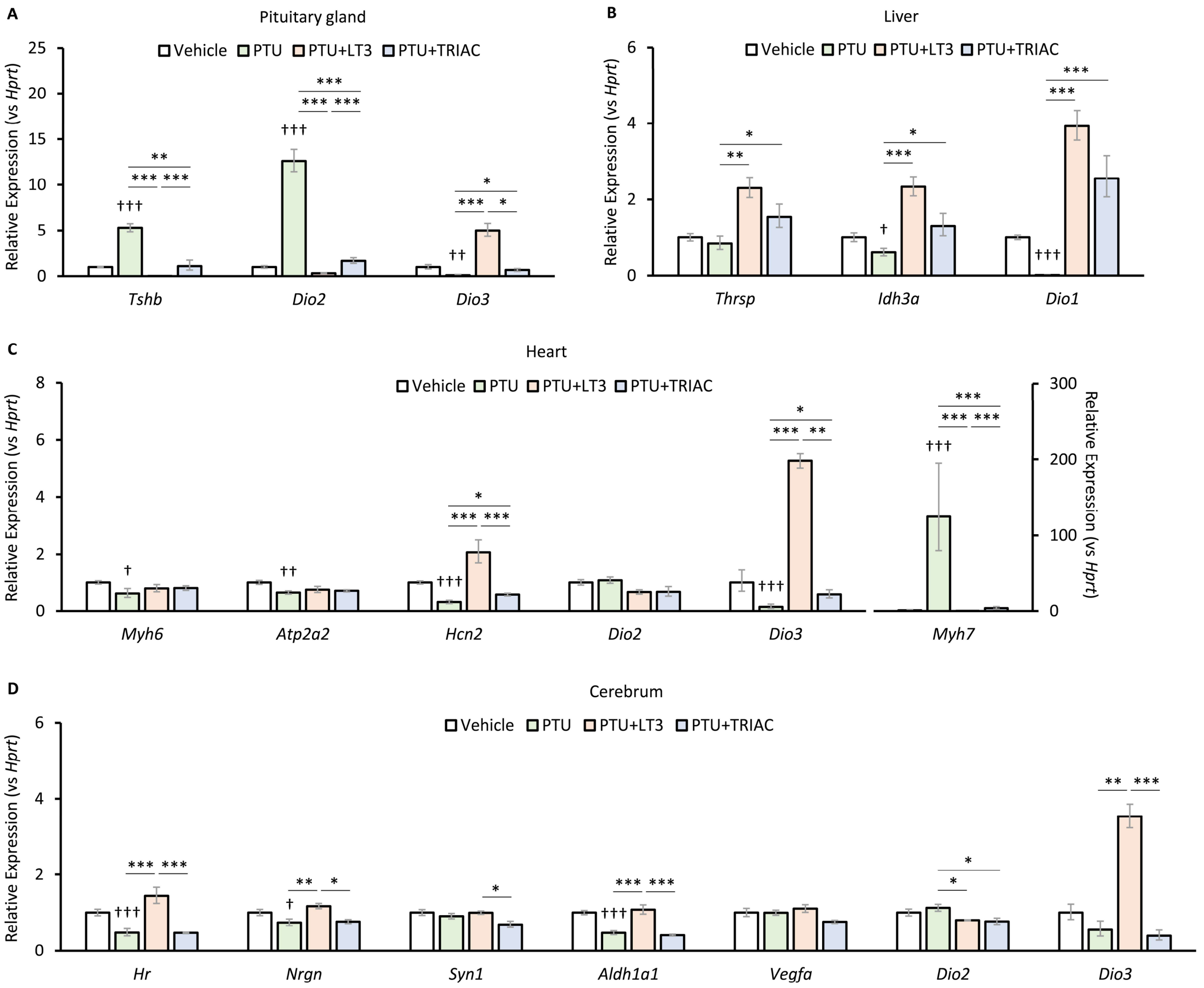


**Supplementary Fig. S2:** Gene expression profiles of hypothyroid mice determined by quantitative RT-PCR normalized using *Hprt* as an internal control. (A) The pituitary gland, (B) the liver, (C) the heart, and (D) the cerebrum. Vehicle, 6-propyl-2-thiouracil (PTU), PTU + TRIAC, $n = 6$ each; PTU + LT3, $n = 5$. Results are shown as fold change versus the vehicle-only control. Statistical analyses were performed by Student's *t*-test for vehicle vs PTU and the *p*-value is presented as †$p < 0.05$, ††$p < 0.01$, and †††$p < 0.001$. ANOVA followed by the Tukey-Kramer test was used for comparisons among PTU, PTU + LT3, and PTU + TRIAC experiments and the *p*-value is presented as *$p < 0.05$, **$p < 0.01$, and ***$p < 0.001$.

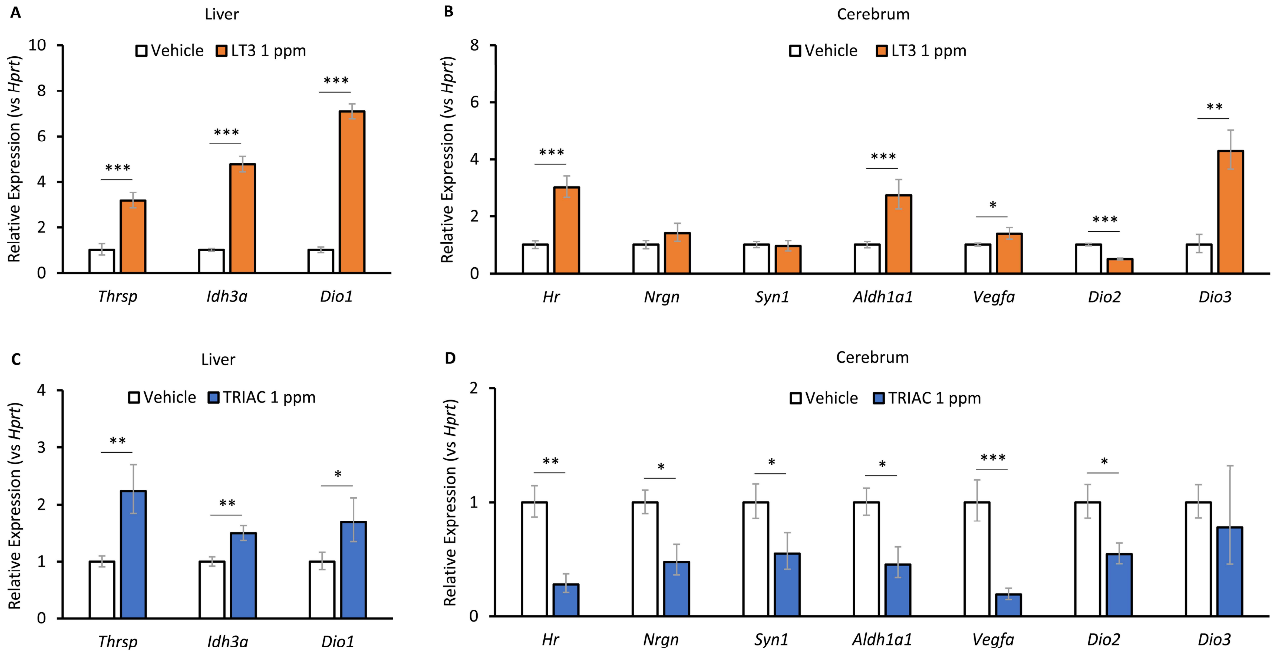


**Supplementary Fig. S3:** Gene expression profiles of euthyroid mice determined by quantitative RT-PCR normalized using *Hprt* as an internal control. (A) The liver and (B) the cerebrum of mice administered with LT3 at 1 ppm ($n = 6$ each). (C) The liver and (D) the cerebrum of mice administered with TRIAC at 1 ppm ($n = 6$ each). Results are shown as fold change versus the vehicle-only group. Statistical analyses were performed by Student's *t*-test. $*p < 0.05$, $**p < 0.01$, and $***p < 0.001$.

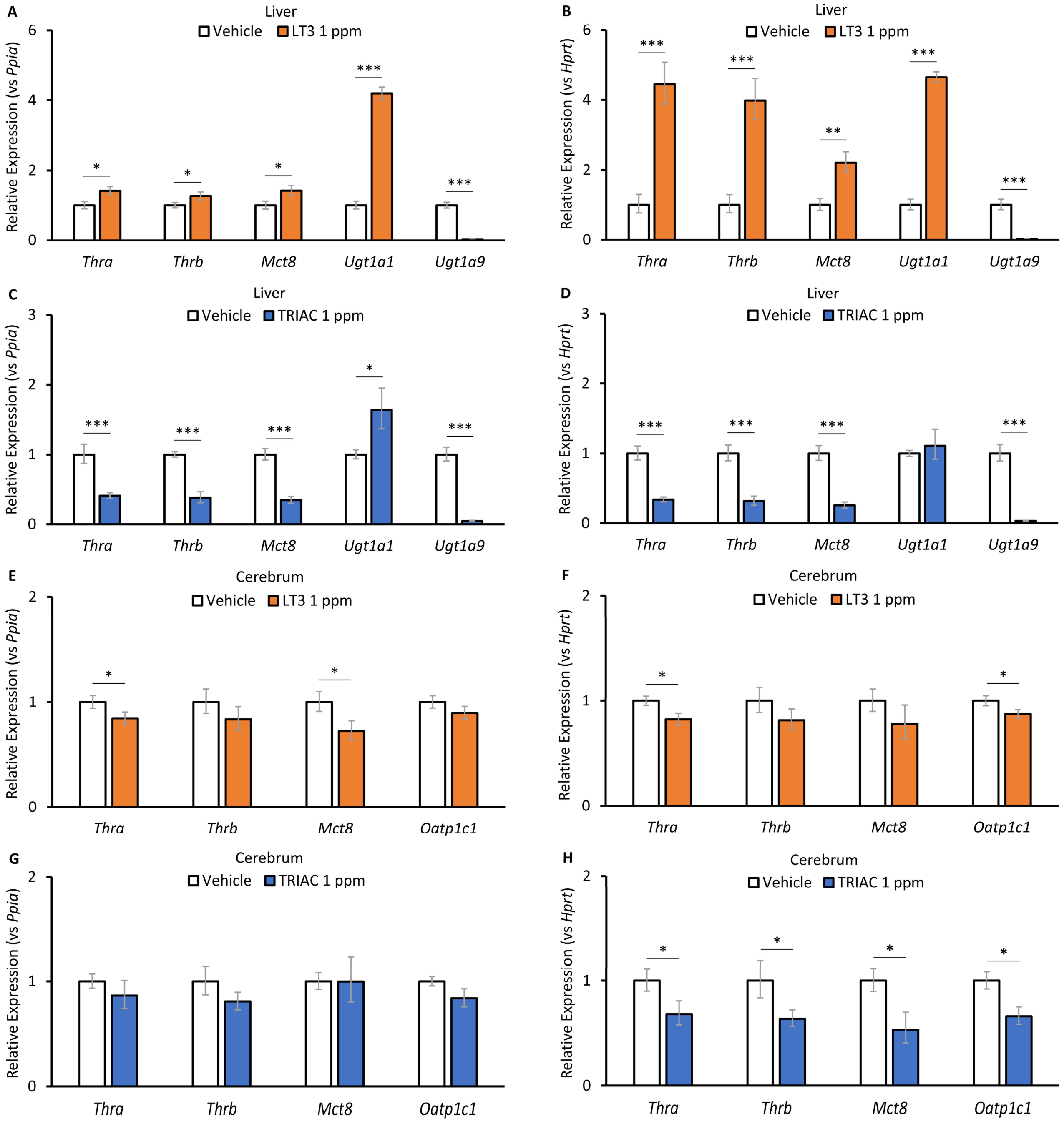


**Supplementary Fig. S4:** Quantitative RT-PCR analyses of euthyroid mice for elements that regulate thyroid hormone actions. (A-D) The liver of mice administered with LT3 at 1 ppm ($n$ = 6 each) (A, B) and that of mice administered with TRIAC at 1 ppm ($n$ = 6 each) (C, D). (E-H) The cerebrum of mice administered with LT3 at 1 ppm ($n$ = 6 each) (E, F) and that of mice administered with TRIAC at 1 ppm ($n$ = 6 each) (G, H). Results are normalized using *Ppia* and *Hprt* as internal controls and shown as fold change versus the vehicle-only group. Statistical analyses were performed by Student's $t$ test. $^*p < 0.05$, $^{**}p < 0.01$, and $^{***}p < 0.001$.

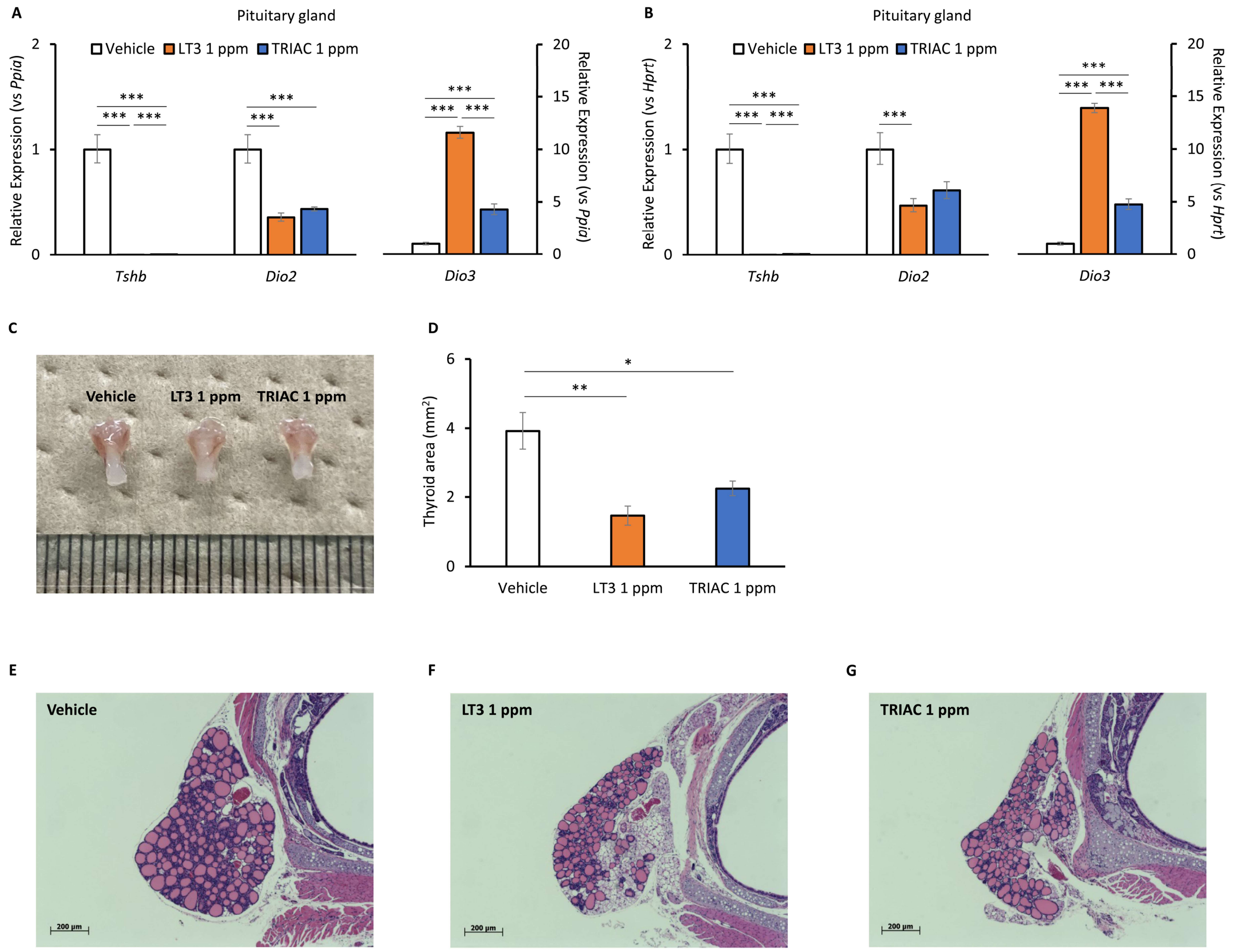


**Supplementary Fig. S5:** Effects of TRIAC and T3 on the hypothalamus-pituitary-thyroid axis in euthyroid mice ($n$ = 4 each). (A, B) Gene expression profiles of the pituitary gland determined by quantitative RT-PCR normalized using *Ppia* (A) and *Hprt* (B) as internal controls. (C) Gross appearance of thyroid glands. The scale on the ruler is 1 mm. (D) Thyroid gland size was measured as thyroid area on photograph. (E-G) Histological images of thyroid glands stained using hematoxylin and eosin. ANOVA followed by the Tukey-Kramer test was used for comparisons and the $p$-value is presented as *$p$ < 0.05, **$p$ < 0.01, and ***$p$ < 0.001.
